# Periplasmic SacB as a robust counter-selection tool for genome engineering in the polyploid bacterium *Zymomonas mobilis*

**DOI:** 10.64898/2026.01.13.699301

**Authors:** Katsuya Fuchino, Christodoulos Astraios, Richard Daniel, James Grimshaw, Manuel Banzhaf, Waldemar Vollmer

## Abstract

The alpha-proteobacterium *Zymomonas mobilis* exhibits exceptional ethanologenic physiology, which makes it a traditional alcoholic beverage producer and a promising chassis for biofuel production. Although genetic tools for this organism have expanded in recent years, a fundamental aspect of its chromosome organization remains to be understood. In particular, *Z. mobilis* has been suggested to exhibit polyploidy, but this feature is not fully confirmed because of discrepancies among studies reporting the copy number of chromosomes. Here, we tagged the chromosome-partitioning protein ParB with a fluorescent marker to visualise its cellular localisation and estimate chromosome copy number in individual cells. Imaging showed that *Z. mobilis* exhibits several distinctive ParB foci throughout the cytoplasm and an accumulated focus at the pole, demonstrating that a single *Z. mobilis* cell contains at >5 copies of the chromosome at the *oriC* regions. After verifying its polyploidy, we sought to establish an efficient counter-selection system, which is crucial for engineering multiple copies of the chromosome. We assessed the efficacy of levan-sucrase (SacB) toxicity in *Z. mobilis*. We found that, despite *Z. mobilis* secreting a native extra-cellular sucrase SacB, heterologous periplasmically-localised *Bacillus subtilis* SacB rendered *Z. mobilis* cells sensitive to sucrose. We successfully used this effect for counter-selection when deleting and inserting targeted DNA sequences into the *Z. mobilis* genome. Together, this work provides important insights and tools for advancing *Z. mobilis* genetics and its biotechnological applications.

**Importance:** *Zymomonas mobilis* is a promising industrial bacterium with capacity to convert sugars into ethanol at nearly maximum theoretical yield. With its expanding use in industrial applications, it is crucial to clarify if individual *Z. mobilis* cells carry multiple copies of chromosome as this has important implications for genome engineering. Two previous studies have used quantitative PCR to address this question, but their reported chromosome copy numbers varied widely from 20 to 100. Here, we used a cell biological approach to estimate the copy number and confirmed that single *Z. mobilis* cell possesses multiple copies. In addition, we show that a SacB-based counter-selection works in *Z. mobilis*, enabling efficient and complete mutation of all chromosome copies.

## Introduction

The alpha-proteobacterium *Zymomonas mobilis* is a natural ethanologen with an exceptional metabolism that can convert sugars into ethanol nearly at the maximal theoretical yield. Its carbon metabolism, the Entner–Doudoroff pathway coupled to pyruvate decarboxylase and alcohol dehydrogenase, is both specific and very rapid (1, 2). These features make *Z. mobilis* one of the most promising chassis organisms for bioethanol production. Beyond ethanol, *Z. mobilis* has recently been engineered to produce various high-value chemicals (3–9).

In efforts to develop *Z. mobilis* for industrial applications, the available genetic tools have expanded considerably in recent years. However, its basic chromosome organisation at the cellular level remains poorly characterised (10). Several studies have suggested that *Z. mobilis* carries multiple copies of its chromosome and is therefore a polyploid species. The first indication of polyploidy came from the transposon mutagenesis study, which revealed insertions in genes considered essential (11). Two other studies using quantitative PCR showed that *Z. mobilis* possesses multiple copies of the chromosome (12, 13). However, the estimated average number of chromosomes per cell varied considerably between reports. Our previous study showed that strains Z6 and Z4 contain approximately 15–20 copies at the replication origin during active growth (12), whereas another study using the same qPCR based method reported that strain Z4 may carry 60 - 100 copies (13).

This significant discrepancy required clarification as the presence of multiple chromosome copies can complicate genetic manipulation. When the applied selection is not tight enough, mixed genotypes are generated, with mutant genomes co-existing with wild-type copies in individual cells. This phenomenon has been reported in previous studies attempting to engineer the *Z. mobilis* genome (14–16).

To address this, we employed a complementary approach to estimate the chromosome copy number: visualising a defined locus using a fluorescent marker and directly counting the signals per single cells (17). This method also offers insights into the spatial organisation of chromosomes (18). Using this approach, we gained further insights in its chromosomal organisation and polyploidy.

Furthermore, having confirmed the polyploidy of *Z. mobilis*, we recognised the need for a robust counter-selection method to enable efficient introduction of desired genetic changes into all chromosomal copies (19). We assessed and successfully applied a SacB-based counter-selection in engineering the *Z. mobilis* genome.

## Material and methods

### Bacterial strains, plasmids, and growth conditions

Bacterial strains and plasmids used in this study are listed in Table S1. *Z. mobilis* was grown in RM complex medium containing glucose (20 g/L), yeast extract (5 g/L), NH_4_SO_4_ (1 g/L), KH_2_PO_4_ (1 g/L), and MgSO_4_ (0.5 g/L). The complex medium was briefly sparged with nitrogen gas to make it micro-aerobic. 12 ml of anaerobic *Z. mobilis* culture were grown at 30°C in a capped 15 ml Falcon tube without shaking. For plotting growth profiles, the optical density was measured in a 96-well plate by SPECTROstar Nano (BMG labtech).

### Construction of *Z. mobilis* strains

For all *Z. mobilis* strains, we used a counter-selection method based on the previously published method (20). Crucially, we used an integrating vector (suicide vector) that carried *Bacillus subtilis sacB* for negative selection (pPK15534 + *SacB^BS^*). To construct the vector, we amplified the *sacB* gene with its native promoter region from *B. subtilis* genomic DNA using Q5 DNA polymerase (New England Biolabs). The product was column purified (QIAquick PCR Purification Kit, Qiagen) and introduced into the suicide plasmid pPK15534 by Gibson assembly (New England Biolabs) (20, 21).

To construct plasmids for gene deletion, the upstream and downstream regions (500 – 1000 bp in size) of the gene of interest, in addition to homologous regions required for Gibson assembly were amplified using Q5 DNA polymerase. For insertion of a tag, the upstream and downstream regions of the insertion sites and the tag were amplified. The obtained DNA fragment was inserted into the linearised plasmid pPK15534 + *sacB^BS^* by Gibson assembly. The sequences of all constructed plasmids were verified by Sanger sequencing (Eurofins Genomics).

Transfer of the suicide vector was performed via conjugation from *E. coli* as described (20). Briefly, the actively growing recipient *Z. mobilis* cells were mixed with *E. coli* WM6026 strain carrying the suicide plasmid, and then plated on RM agar medium with 0.1 mM meso-diaminopimelate (DAP) and 2% glucose overnight at 30 °C. The next day, the grown cells were scraped and resuspended in liquid RM + 2% glucose, followed by an incubation at 30°C for 2 h. Cultures were then diluted (typically 10x and 100x, depending on the density) into phosphate-buffered saline (PBS) and 100 μL of each dilution was spread on RM + 2% glucose agar medium with 90 μg/mL chloramphenicol to select for the cells that received and integrated the plasmids.

The colonies that grew on the selection plates were then checked for the presence of plasmid in their chromosomal DNA by colony PCR. Verified colonies were then grown in liquid RM + 2% glucose without selection overnight (excision of the plasmids). Next day, the cells were diluted in PBS (10, 100 and 1000x), and 100 μL from each diluted culture were spread on RM agar supplemented with sucrose (initially using 3%, 4% or 5%) and the plates were incubated for two days. We found that 3% sucrose provided tight selection and used this concentration for generating most of the strains in Table 1, but 5% sucrose was also effective. After the colonies appeared, single colonies were streaked on RM with 2% glucose, then colony PCR was performed to identify colonies with the desired genotype. The genomic DNA of several generated strains were extracted using a genomic DNA kit (Sigma), sequenced by whole genome sequencing (MicrobesNG) and analysed by CLC Genomics Workbench (Qiagen). Oligonucleotides used in this study are listed in Table S2.

**Table 1.** The summary of outcome of genotype after the excision of the plasmid integrated into the wild-type strain Z6.

| Introduced mutation/tag | number of tested colonies | Genotype |  |  |
| --- | --- | --- | --- | --- |
|  |  | mixed | wild-type | targeted |
| $\Delta Z6\_1449$ | 3 | 1 | 1 | 1 |
| $Z6\_1449$-His x 6-FLAG | 6 | 0 | 4 | 2 |
| $Z6\_0812$-mCherry | 10 | 0 | 8 | 2 |
| $\Delta Z6\_1449$ C-ter truncation | 11 | 0 | 10 | 1 |
| $\Delta Z6\_0810$ | 6 | 0 | 5 | 1 |
| $\Delta Z6\_0149$ | 12 | 1 | 9 | 2 |
| $\Delta Z6\_1271$ | 22 | 0 | 15 | 7 |
| $\Delta Z6\_1369$ | 24 | 0 | 15 | 9 |
| $Z6\_1203\text{-GFPsf}$ | 28 | 0 | 6 | 22 |
| $\Delta Z6\_1479$ | 28 | 0 | 26 | 2 |
| $\Delta Z6\_0234$ | 28 | 0 | 24 | 4 |

## Microscopy analysis

A culture of growing *Z. mobilis* (0.4 µl) was spotted onto a 1% agarose-pad made of RM growth medium and covered by a small cover-glass. Phase contrast and fluorescence images of *Z. mobilis* cells were taken using a Nikon Eclipse Ti microscope (Nikon) equipped with a Plan Apochromat 100x objective with an NA of 1.40 and Ph3 Phase plate (Nikon) and a Prime sCMOS camera (Teledyne Photometrics). ParB-sfGFP and FM 4-64 were imaged using 49002 and 49008 filter cubes respectively (Chroma). Images were acquired using Nikon NIS elements AR software. Image analysis was performed using MicrobeJ (22). For chromosome foci counting, we used Maxima function in MicrobeJ using tolerance value as 850. For staining membrane, the cells were incubated with FM4-64 (2 μg/mL), for 5 minutes prior to imaging.

## Results

### ParB-sfGFP forms multiple foci in single *Z. mobilis* cells

To visualise the *oriC* site of chromosome in *Z. mobilis* cells, we expressed chromosome partitioning protein ParB fused to the fluorescent protein sfGFP (super-folder variant of GFP) (23). ParB has been shown to play a central role in segregating of replicating chromosomes in several bacteria, assembling at *parS* DNA sites located near the origins of replication (*oriC*) (24). Bioinformatic analysis predicted four *parS* sites near the *oriC* in the *Z. mobilis* chromosome (25). Thus, ParB sub-cellular localisation should serve as a marker for counting chromosomal origins in a single *Z. mobilis* cell. (17) We generated a *Z. mobilis* strain in which *parB*-*sfGFP* was introduced at the original locus (*Z6_1203*) encoding ParB. The strain did not show any growth or morphological defects (Fig. S1), indicating that the ParB-sfGFP is a functional fusion. The fluorescence imaging of growing Z6 ParB-sfGFP cells showed that the ParB-sfGFP exhibited several sub-cellular localisation patterns (Fig. 1AB): 1) Multiple distinctive sfGFP foci were distributed throughout the cytoplasm within a cell with noticeable variation in intensity. Interestingly, the *Z. mobilis* ParB-foci did not exhibit an ordered spatial arrangement, in contrast to the organized ParB positioning reported for several other bacteria including polyploid species (17, 26–28). 2) There were often smeared signals around the foci, which might suggest the spreading of ParB (29). 3) A larger focus with an intense signal was observed at one of the cell poles. The polar focus exhibited, on average, more than two-fold higher fluorescence intensity compared with non-polar foci. (Fig. 1C). 4) When the cells were washed with PBS, the fluorescent foci disappeared and the signal became diffused throughout the cytoplasm (Fig. S2), indicating that the foci were unlikely to be inclusion bodies.

**Fig. 1.**
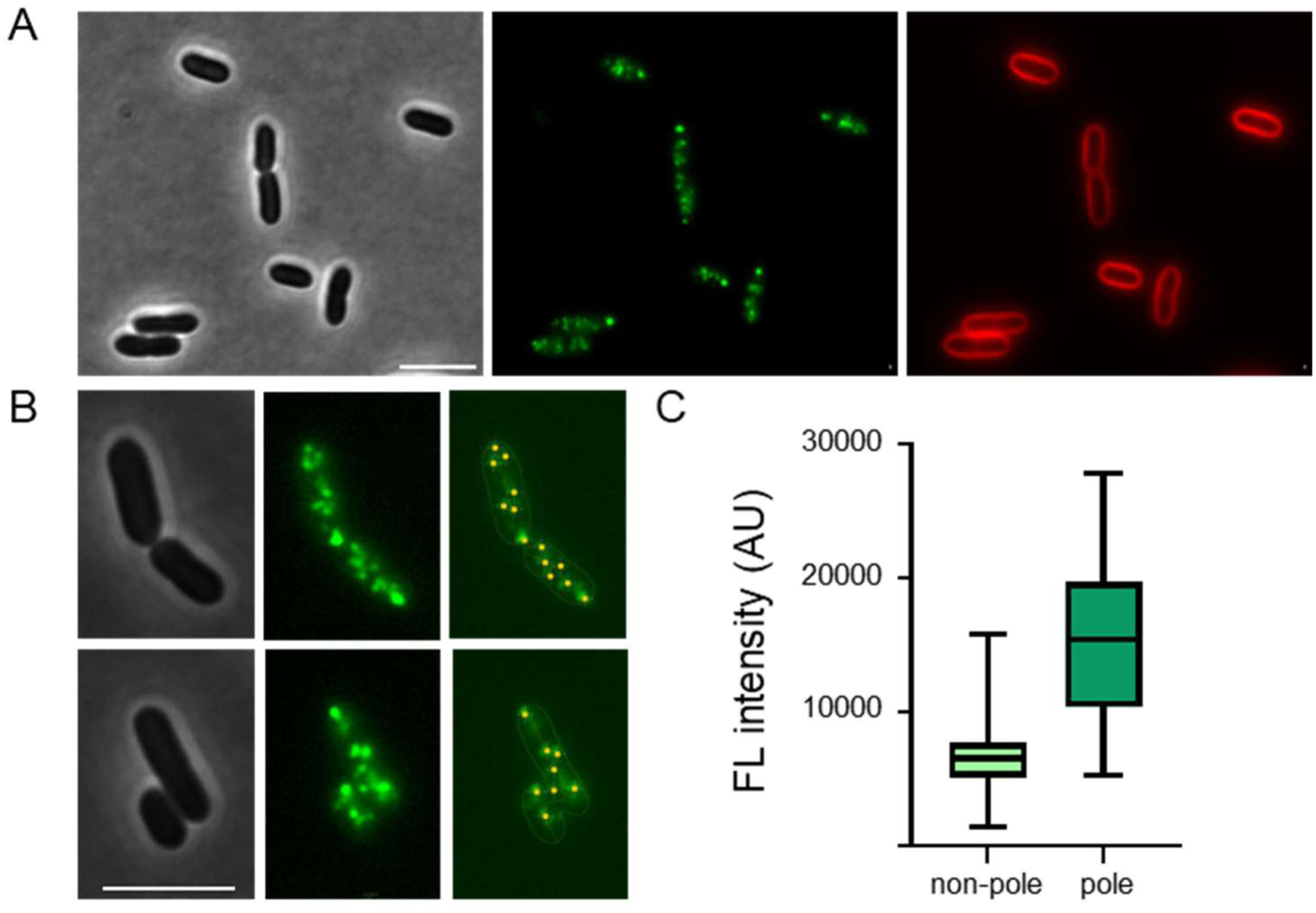
ParB-sfGFP sub-cellular localisation in *Z. mobilis*. (A) Phase contrast (left) and fluorescent images (middle, right) of growing Z6 ParB-sfGFP cells under standard growth conditions. For visualising membrane, the cells were stained with FM4-64 (right panel). Scale bar, 5 μm. (B) ParB-sfGFP detection in the *Z. mobilis* cells using microbeJ (22). Distinctive foci were detected from the fluorescence images. Scale bar, 5 μm. (C) Box-and-whisker plot presenting ParB-sfGFP fluorescence intensity of polar focus and foci detected from other cell regions. Biological replicates N = 3, total measured foci N = 1063.

We counted the distinctive fluorescent foci within cells using microbeJ (22) which enables automated detection and quantitation of foci-signals within cells. However, the number of detectable foci likely represents a lower-bound estimate of the *oriC* numbers as *Z. mobilis* chromosomes appear to lack ordered spatial organization within the limited cytoplasmic space, which could lead to co-localization of multiple ParB-sfGFP foci (see Discussion). Despite this potential technical limitation, we estimated a minimum chromosomal copy number of 5.6 +-1.8 based on the separated ParB-sfGFP foci (from evaluating 561 cells).

### Periplasmic Levan sucrase is toxic to *Z. mobilis* cells in the presence of sucrose

Having demonstrated that individual *Z. mobilis* cell contains multiple copies of chromosome, we recognised that engineering the *Z. mobilis* genome requires an efficient counter-selection system. In polyploid bacteria, the presence of multiple chromosome copies could result in heterogeneous genotype during genome editing, with wild-type and mutated genes coexisting on different chromosomes within the same cell. Thus, positive selection strategies, such as antibiotic resistance markers, might lack sufficient selection, because it allows the survival of cells that retained unmodified, wild-type chromosome copies. In contrast, a negative selection eliminates cells carrying unmodified chromosomes, efficiently selecting cells in which all chromosomal copies altered.

Currently, the widely used method to delete gene in *Z. mobilis* is homologous recombination-based method utilizing a suicide vector carries flanking region of the targeted gene (30), particularly the method developed by Ral *et al.* which relies on antibiotic resistance for the integration of the plasmid, and the loss of a GFP fluorescence reporter for the excision of plasmid from the chromosome (20). This powerful method has been previously used by us to successfully generate mutant strains (31). However, we found that the last selection (excision of the integrated plasmid) was not efficient, which resulted in obtaining mixture of cells that underwent the excision and cells carrying the integrated plasmid. This was also observed in another study that used the method (14). The high rate of false positive cells is presumably due to the presence of multiple chromosome copies. In addition, the method requires using time-consuming Fluorescence-Activated Cell Sorting (FACS). Thus, we aimed to develop an alternative, practical negative selection method that forces the excision event and allows simple isolation of cells that have the desired genetic manipulation.

Although the *sacB* encoding levan sucrase was previously used as a counter-selection in *Z. mobilis* (32) the use of levan sucrase was later questioned by the *Z. mobilis* research community because the *Z. mobilis* genome encodes its own *sacB* gene and can grow on sucrose as a sole carbon source. However, the native *Z. mobilis* SacB was reported to be extracellular (33) while the toxicity of levan sucrase manifests within the periplasm in Gram-negative bacteria (34). SignalP analysis (35) predicts a periplasmic localisation for *Bacillus subtilis* SacB in Gram-negative bacterial cells, but not for the *Z. mobilis* SacB consistent with the previous report (33).

We next examined whether expressing the periplasmic SacB^BS^ (*Bacillus subtilis* SacB) could be toxic to *Z. mobilis* cells in the presence of sucrose and thus could potentially serve as a counter-selector as in other Gram-negative bacteria. We generated the *Z. mobilis* strain carrying *sacB^BS^* inserted into the chromosome at the adjacent region of Z6_1449 (31), and tested growth in RM complex medium supplemented with either 2% glucose or 5% sucrose. While the strain grew as wild-type in the RM agar medium supplemented with 2% glucose, it failed to grow in the presence of 5% sucrose (Fig. 2). This demonstrates the SacB-toxicity depends on the cellular localisation of SacB and indicates that periplasmic SacB^BS^ is applicable for negative selection in *Z. mobilis*.

**Fig. 2.**
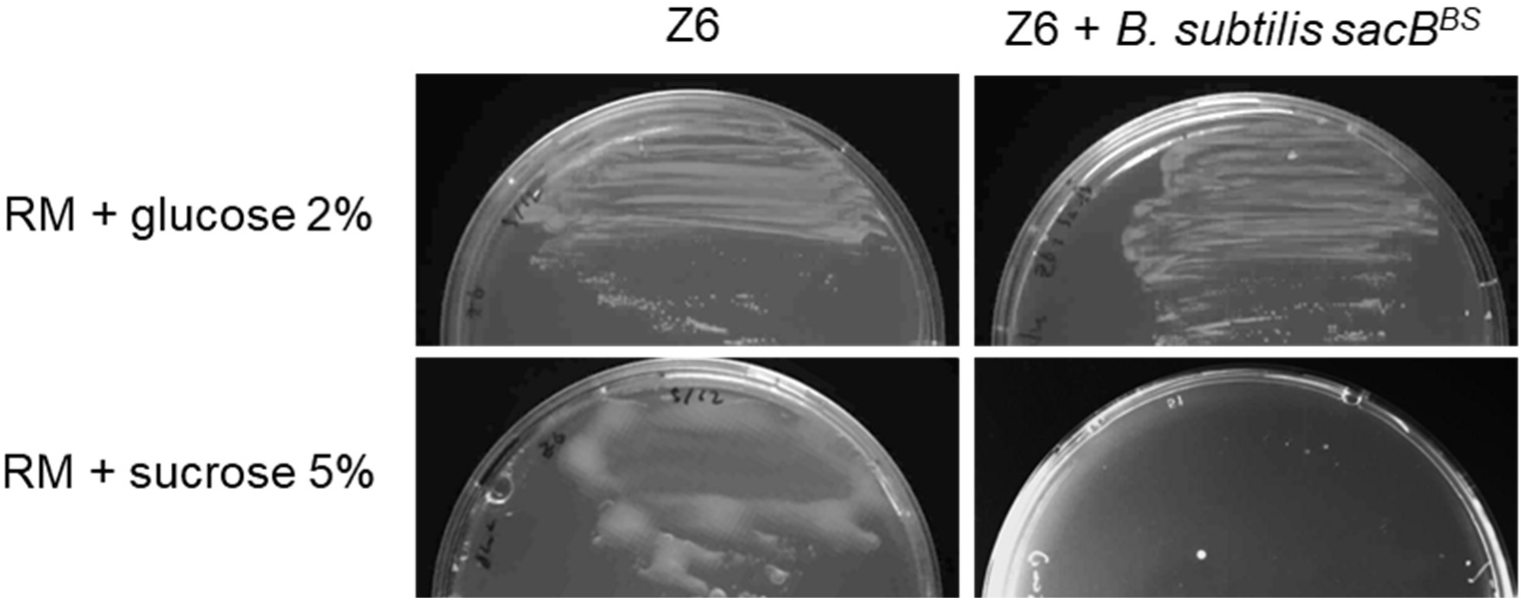
Periplasmic SacB^BS^ prevents growth of *Z. mobilis* cells in the presence of sucrose. (top) Both Z6 and Z6 + SacB^BS^ strains grew on RM supplemented with 2% glucose. (bottom) Z6 exhibited lysing colonies on RM + sucrose 5% (left side) while Z6 + SacB^BS^ strain did not grow at all in the presence of 5% sucrose (right side).

### Sucrose in the growth medium triggers explosive lysis of wild-type *Z. mobilis* cell

Close inspection of the wild-type strain grown on RM agar-medium supplemented with sucrose indicated that the colonies displayed partial lysis (Fig. 2). The lysis is likely due to the activity of its native, extracellularly secreted SacB producing toxic levan polymer. Such stress-induced lysis might be a problem for a counter-selection system, as continuous stress might increase the likelihood of unwanted, secondary mutations during genome engineering. To observe the lysis at a cellular level, we performed time-lapse imaging on the wild-type strain Z6 growing on glucose or sucrose in the complex medium. While the majority of cells fed with sucrose grew well, we observed abrupt lysis of a few single cells during imaging (Fig. 3). The cytoplasmic content released from the cell pushed other cells on the agar-pad within 10 minutes during the imaging (Fig. 4, bottom panels). The explosive lysis was observed in other time-lapse videos, and typically affected only one cell within a micro colony consisting of few hundred cells. This low level of lysis of Z6 in the presence of sucrose presumably explains the slimy appearance of cells on the RM agar-medium with sucrose (Fig. 2 left bottom) and the majority of cells remained viable and continued to actively grow. Consistent with this, *Z. mobilis* cultures grown in liquid medium containing sucrose exhibited no apparent growth defects as previously reported (36, 37).

**Fig. 3.**
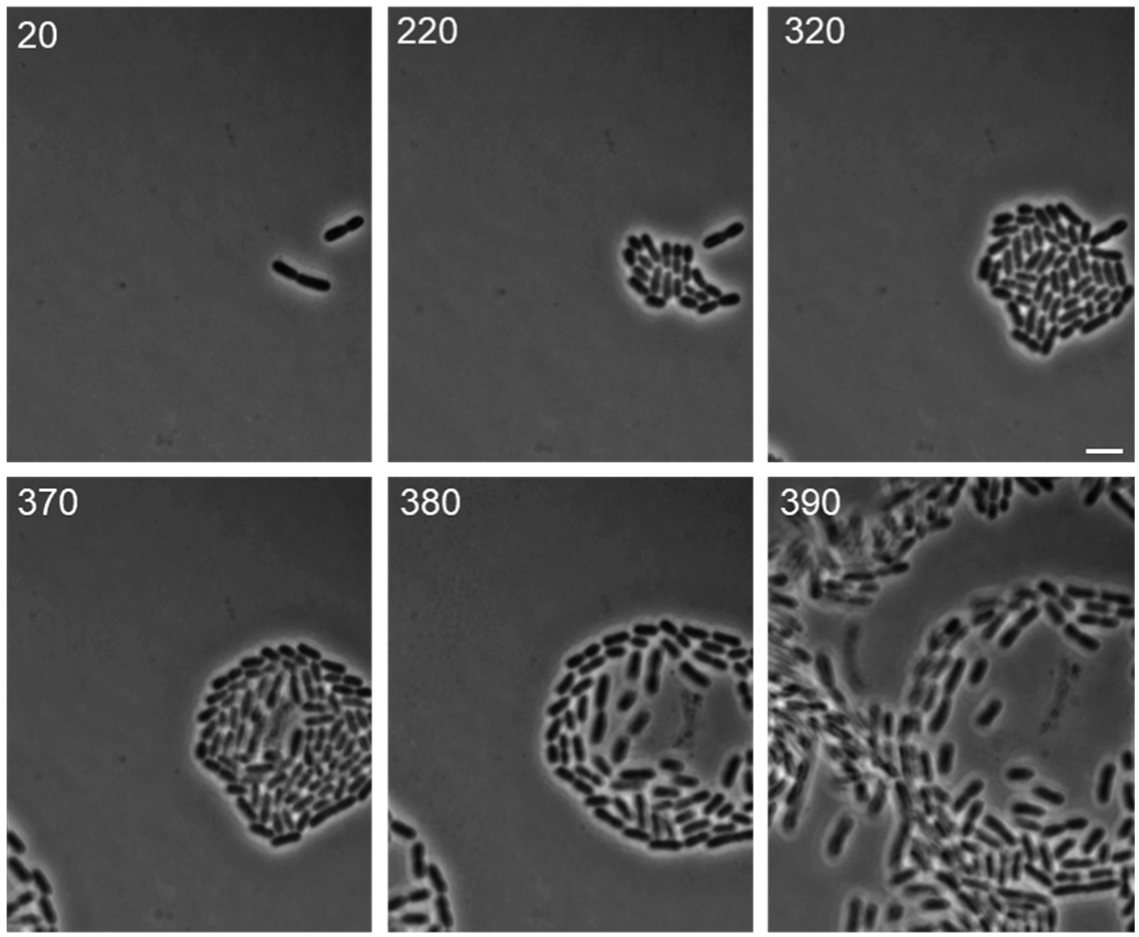
*Z. mobilis* cells growing in the presence of 5% sucrose. Images show that one cell exhibits explosive lysis, releasing its cytoplasm content that washes away other cells on RM medium. (Time point from top left to bottom right, 20, 220, 320, 370, 380, 390 minutes after the imaging started) Scale bar, 5 μm.

### Periplasmic-SacB^BS^ counter selection is applicable in engineering the *Z. mobilis* genome

Next, we applied the SacB^BS^-based counter-selection system by integrating it into the previously published method (20) by replacing the plasmid-excision step based on FACS with sucrose-sensitivity selection. In this system, cells retaining the integrated plasmid are killed in the presence of sucrose. Consequently, only cells in which the plasmid has been excised from all chromosomal copies survive, ensuring complete plasmid removal. Using this modified method, we readily obtained colonies that appeared to have lost the targeted gene, as checked by PCR. To verify the absence of wild-type gene in all chromosomes, we also analysed the whole genome sequence of three independent strains (Δ*Z6_1449*, Δ*Z6_0810* and Δ*Z6_1479*). The analysis confirmed the 100%, homozygous deletion of targeted gene without the wild-type copy, and no additional mutations in other regions of the chromosomes. We further utilised the SacB counter-selection system to introduce gene fusions or tags into coding sequences (epitope or fluorescent tag) into the chromosomes. We successfully obtained the strains with desired tags at the precise location in all copies of chromosome, confirmed by whole genome sequencing (Z6_1449-His x 6-FLAG). Table 1 summarises the generated strains and frequencies of obtaining the desired genotype. On average, 29.8% of the mutant colonies (based on PCR analysis) were obtained from the tested colonies by the sucrose selection, which we found sufficiently effective and practical. Taken together, these results demonstrate that the SacB-based counter selection is a reliable and efficient tool for engineering the *Z. mobilis* genome.

## Discussion

This study adds new insights into the polyploidy and chromosomal organization of *Z. mobilis*. Our initial objective was to determine the chromosome copy number at the single-cell level using fluorescently tagged ParB that binds to *parS* site in the chromosome. However, this proved to be challenging due to the spatial organisation of chromosomes within the cell, and presumably led to underestimating the copy numbers because clustered foci were counted as single signal. In particular, the intense polar focus likely represents multiple *oriC* sites. In addition, *Z. mobilis* cells are approximately 1.1 - 1.4 µm wide, hence foci positioned above or below the focal plane might be missed when imaging of a single-plane. We aimed to visualise 3-dimentional imaging of the signals by Z-stacking of fluorescence imaging, but the foci were found to be dynamic, which made counting foci from stacked images unreliable (data not shown). Consequently, the copy numbers obtained from the imaging likely represent a lower-bound estimate. Consistent with this, the copies numbers obtained from the sfGFP signals were lower than those previously measured using quantitative PCR method (12).

The intense polar localization of ParB has been reported for other alpha-proteobacteria (28, 38) and might reflect a conserved feature of chromosome replication in this bacterial group, indicating that the observed intense foci might represent the ongoing replicating site.

The physiological benefits of polyploidy are not yet understood in *Z. mobilis*. Previous work showed that the strain Z4 can grow under phosphorus-depleted conditions, presumably by using its chromosome as a phosphorus source, which might be a strategic advantage of being polyploid (39). Another proposed advantage of polyploidy is to enhance genomic stability. A study on long-term co-cultivation of *Z. mobilis* with *Saccharomyces cerevisiae* to induce genetic changes observed no major mutations in the *Z. mobilis* chromosome, potentially due to the presence of multiple chromosome copies that could facilitate reversion of mutations (40). On the other hand, several studies have employed adaptive evolution to obtain *Z. mobilis* strains with desirable traits (41–43) and as it remains unclear in some studies whether the resulting populations were genetically homogeneous or contained mixed genotypes, further examination in the context of the polyploid nature of *Z. mobilis* might be needed.

In addition to advancing our understanding of chromosome organization, this study also provides an additional genetic tool for genome engineering in *Z. mobilis*. We showed that the frequently-used SacB counter selection is applicable to *Z. mobilis*, and we implemented in the existing homologous recombination based method, showing high frequency (29.8%) of obtaining the desired genotype. Recently, two different counter-selection systems for *Z. mobilis* have been reported (44, 45). The availability of multiple robust counter selection strategies should be advantageous for genome engineering in *Z. mobilis*. Efficient counter-selection is not only useful for homologous recombination–based approaches but also for recently developed CRISPR-based systems, which needs reliable plasmid curing (46).

## Acknowledgements

We thank Prof. Patricia J. Kiley (University of Wisconsin–Madison) for donating the plasmid pPK15534. This work was supported by Newton International Fellowship (The Royal Society) to K.F. (NIF\R1\221190)

