## Supplemental Figure S1-S2 and Supplemental table S1 -2 for "Periplasmic SacB as a robust counter-selection tool for genome engineering in the polyploid bacterium *Zymomonas mobilis*"

### Supplementary figures

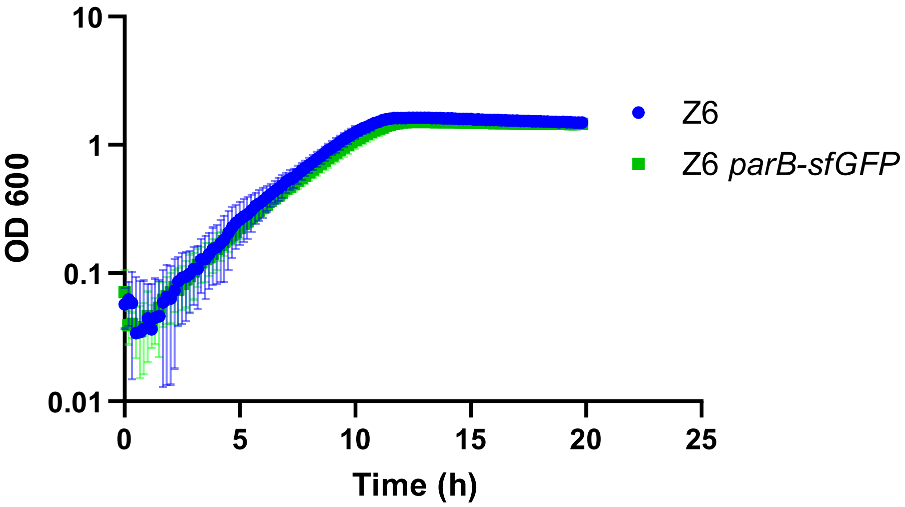


Fig. S1. Growth profiles of *Z. mobilis* strains Z6 (wild-type) and Z6 + *parB-sfGFP* under regular growth conditions. Biological replicates N = 3. The error bars represent the standard deviation.

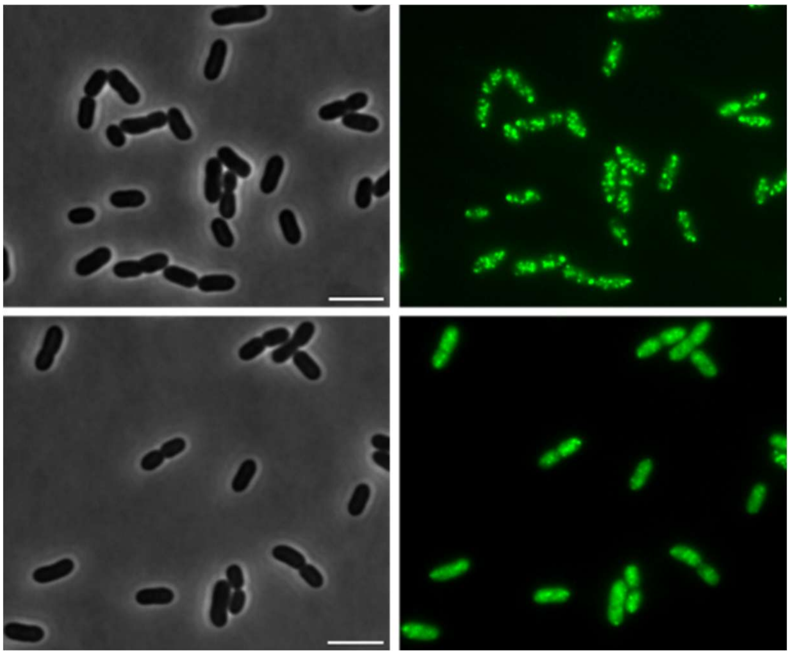

Fig. S2 ParB-sfGFP localisation in the strain Z6 + *parB-sfGFP* washed by PBS prior to imaging. The phase contrast (left side) and fluorescence (right side) image of the *Z. mobilis* ParB-sfGFP cells (top) directly imaged from the culture and (bottom) being washed by PBS before mounting on the agar-pad. The washed cells have a dispersed sfGFP signal, indicating that the stress caused dissociation of the ParB-sfGFP from the chromosomes. Scale bar, 5 µm.

### Supplementary tables

Table S1. Bacterial strains and plasmids used in this study.

| Bacterial strains | Genotype/ description/reference |
|---|---|
| *Zymomonas mobilis* Z6 | ATCC29191 purchased from DSMZ |
| *Escherichia coli* Dh5a | Cloning strain (Lab stock) |
| *Escherichia coli* WM6026 | Conjugation strain (1) |
| *Zymomonas mobilis* + SacB | *B. subtilis* sacB at *Z6_1449* (this study) |
| *Zymomonas mobilis ΔZ6_1449* | *ΔZ6_1449* (this study) |
| *Zymomonas mobilis Z6_1449-His x 6-FLAG* | *Z6_1449-His x 6-FLAG* (this study) |
| *Zymomonas mobilis Z6_0812-mCherry* | *Z6_0812-mCherry* (this study) |
| *Zymomonas mobilis ΔZ6_1449 C-ter* | *ΔZ6_1449 C-terminus truncation* (this study) |
| *Zymomonas mobilis ΔZ6_0810* | *ΔZ6_0810 (this study)* |
| *Zymomonas mobilis ΔZ6_0149* | *ΔZ6_0149* (this study) |
| *Zymomonas mobilis ΔZ6_1271* | *ΔZ6_1271* (this study) |
| *Zymomonas mobilis ΔZ6_1369* | *ΔZ6_1369* (this study) |
| *Zymomonas mobilis Z6_1203-sfGFP (parB-sfGFP)* | *Z6_1203-sfGFP* (this study) |
| *Zymomonas mobilis ΔZ6_1479* | *ΔZ6_1479* (this study) |
| *Zymomonas mobilis ΔZ6_0234* | *ΔZ6_0234* (this study) |

| Plasmids | Description/reference |
|---|---|
| pPK15534 | Suicide vector (1) |
| pPK15534 + *sacB* | pPK15534 carrying *B. subtilis sacB* (this study) |
| pPK15534 + *sacB Δ1449* | pPK15534 + *sacB* carrying *Δ1449* cassettes (this study) |
| pPK15534 + *sacB Z6_1449-His x 6-FLAG* | pPK15534 + *sacB* carrying *Hisx6-FLAG* (this study) |
| pPK15534 + *sacB Z6_0812-mCherry* | pPK15534 + *sacB* carrying mCherry (this study) |
| pPK15534 + *sacB ΔZ6_1449 C-ter* | pPK15534 + *sacB* carrying *ΔZ6_1449* C-ter (this study) |
| pPK15534 + *sacB ΔZ6_0810* | pPK15534 + *sacB* carrying *ΔZ6_0810* cassettes (this study) |
| pPK15534 + *sacB ΔZ6_0149* | pPK15534 + *sacB* carrying *ΔZ6_0149* cassettes (this study) |
| pPK15534 + *sacB ΔZ6_1271* | pPK15534 + *sacB* carrying *ΔZ6_1271* cassettes (this study) |
| pPK15534 + *sacB ΔZ6_1369* | pPK15534 + *sacB* carrying *ΔZ6_1369* cassettes (this study) |
| pPK15534 + *sacB Z6_1203-sfGFP* | pPK15534 + *sacB* carrying *Z6_1203-sfGFP* (this study) |
| pPK15534 + *sacB ΔZ6_1479* | pPK15534 + *sacB* carrying *ΔZ6_1479* cassettes (this study) |
| pPK15534 + *sacB ΔZ6_0234* | pPK15534 + *sacB* carrying *ΔZ6_0234* cassettes (this study) |

Reference (1): Lal *et al*, Front. Microbiol. 10:2216. doi: 10.3389/fmicb.2019.02216

Table S2. Oligonucleotides used in this study.

| Oligo | Sequence | Use |
|---|---|---|
| **NKF1** | TTTTTTCAACCACAAGATAACGTATGGGGCTGACTTCAGGTGC | ***ΔZ6_1449*** |
| **NKF2** | TGTTCAATGCCATCATGAATGTAATTCTCATGTTTGACAGCTTATCAC | |
| **NKF3** | CCTGAAGTCAGCCCCATACGTTATCTTGTGGTTGAAAAAAATACCG | |
| **NKF4** | ATTATCCCGAATTGTATTTTCAGTCTTTACTTTGTCCATCCATGACAG | |
| **NKF5** | GATGGACAAAGTAAAGACTGAAAATACAATTCGGGATAATAAGGATCGG | |
| **NKF6** | CTGTCAAACATGAGAATTACATTCATGATGGCATTGAACATTTCTTGGG | |
| **NKF47** | AAGCCGAAGCCGATCCACCATCGTATGGGGCTGACTTCAGGTGC | ***Z6_1449-Hisx6-FLAG*** |
| **NKF48** | CACCTGAAGTCAGCCCCATACGATGGTGGATCGGCTTCGGCTTCGG | |
| **NKF49** | TTAGTGGTGGTGGTGGTGGTGCTTGTCGTCGTCGTCCTTGTAGTCTTCCCGACCCGATGGGGTGG | |
| **NKF50** | CAAGGACGACGACGACAAGCACCACCACCACCACCACTAAAAAATACAATTCGGGATAATAAGGATCGG | |
| **NKF51** | GTGATGGGTTAAAAAGGATCTTGGCGAAAATGAGACGTTGATCGGC | ***sacB*** **insertion into pKK15534** |
| **NKF52** | AATTAACAGTTAACAAATAACCAGGGCTTCCCGGTATCAACAGGG | |
| **NKF53** | TCAACGTCTCATTTTCGCCAAGATCCTTTTTAACCCATCACATATACCTGC | |
| **NKF54** | TTGATACCGGGAAGCCCTGGTTATTTGTTAACTGTTAATTGTCCTTGTTC | |
| **NKF78** | TATCTTTCATCGGGCTGATGCGTATGGGGCTGACTTCAGGTGC | ***ΔZ6_1449*** **C-terminus truncation** |
| **NKF79** | CCTGAAGTCAGCCCCATACGCATCAGCCCGATGAAAGATACGCC | |
| **NKF80** | ATTATCCCGAATTGTATTTTCTAGATCGCCCGGTTTTGAAAAATAACGC | |
| **NKF81** | TTCAAAACCGGGCGATCTAGAAAATACAATTCGGGATAATAAGGATCGG | |
| **NKF91** | AAATGCCCCGCAACGATTGCCGTATGGGGCTGACTTCAGGTGC | **Z6*_0812-mCherry*** |
| **NKF92** | TGCGCTACTTCAATCGCTTTGTAATTCTCATGTTTGACAGCTTATCAC | |
| **NKF93** | ACCTGAAGTCAGCCCCATACGGCAATCGTTGCGGGGCATTTAGAAC | |
| **NKF94** | TCCTCCTCGCCCTTGCTCACAAACTGGTAATAAATAAAGGGCGCGAC | |
| **NKF95** | GCCCTTTATTTATTACCAGTTTGTGAGCAAGGGCGAGGAGGATAAC | |
| **NKF96** | CATAATGCCTTATTCCCCTACTACTTGTACAGCTCGTCCATGCCG | |
| **NKF97** | TGGACGAGCTGTACAAGTAGTAGGGGAATAAGGCATTATGCCTAATGAC | |
| **NKF98** | CTGTCAAACATGAGAATTACAAAGCGATTGAAGTAGCGCATGAAG | |
| **NKF59** | TTTATCCGAGACGTTATCTGCGTATGGGGCTGACTTCAGGTGC | **ΔZ6_0810** |
| **NKF60** | AAAACGCTCACAGGCAAGAAGTAATTCTCATGTTTGACAGCTTATCAC | |
| **NKF61** | CCTGAAGTCAGCCCCATACGCAGATAACGTCTCGGATAAAGAGAAG | |
| **NKF62** | AATAGCCTCCCATATTAAAACGACAACTCCATAACCTGCGC | |
| **NKF63** | CGCAGGTTATGGAGTTGTCGTTTTAATATGGGAGGCTATTCAGCCTC | |
| **NKF64** | CTGTCAAACATGAGAATTACTTCTTGCCTGTGAGCGTTTTTGGAG | |
| **NKF133** | CAAGTGTAGAATTGTCAGGACGATCGTATGGGGCTGACTTCAGGTGC | ***ΔZ6_0149*** |
| **NKF134** | GATTAGAAGCCAATAAAGATGTAATTCTCATGTTTGACAGCTTATCAC | |
| **NKF135** | GCACCTGAAGTCAGCCCCATACGATCGTCCTGACAATTCTACACTTGTCTCC | |
| **NKF136** | TTTTAAGGCGCAGCTAAATCAGAGGAACTGATAAAACAGACTAAAGTTTCTTGCC | |
| **NKF137** | ACTTTAGTCTGTTTTATCAGTTCCTCTGATTTAGCTGCGCCTTAAAAATAG | |
| **NKF138** | AAGCTGTCAAACATGAGAATTACATCTTTATTGGCTTCTAATCCTTCTGAAAAAGG | |
| **NKF169** | GCTTCGCTTTTTAGCAAGGCGCCGTATGGGGCTGACTTCAGGTGC | ***ΔZ6_1271*** |
| **NKF170** | ACTTTCAGGGTCAAGAAACCGTAATTCTCATGTTTGACAGCTTATCAC | |
| **NKF171** | CACCTGAAGTCAGCCCCATACGGCGCCTTGCTAAAAAGCGAAGCAG | |
| **NKF172** | AGCCGTAACCGGTTTGAAAAATCAATCATGCTCACAATGTGCCTG | |
| **NKF173** | GCACATTGTGAGCATGATTGATTTTTCAAACCGGTTACGGCTTTTTAC | |

| | | |
|---|---|---|
| **NKF174** | GCTGTCAAACATGAGAATTACGGTTTCTTGACCCTGAAAGTTTGACC | |
| **NKF198** | AACATCGCCGCCCATCCACACCCGTATGGGGCTGACTTCAGGTGC | ***ΔZ6_1369*** |
| **NKF199** | GCGGACACTATGAACCAGAAGTAATTCTCATGTTTGACAGCTTATCAC | |
| **NKF200** | CCTGAAGTCAGCCCCATACGGGTGTGGATGGGCGGCGATGTTC | |
| **NKF201** | AAAACGGATTTTGAGAAGTTTAAAAGAGCCTACCTTTCTTATTACTTCAGC | |
| **NKF202** | AGTAATAAGAAAGGTAGGCTCTTTTAAACTTCTCAAAATCCGTTTTAAGCGG | |
| **NKF203** | AAGCTGTCAAACATGAGAATTACTTCTGGTTCATAGTGTCCGCCGC | |
| **NKF257** | TGATAGGCCTTGGCTTCCTCAACGTATGGGGCTGACTTCAGGTGC | ***Z6_1203*-sf*GFP*** |
| **NKF258** | CGCTCGACAAGCTGGGTGGTGTAATTCTCATGTTTGACAGCTTATCAC | |
| **NKF259** | ACCTGAAGTCAGCCCCATACGTTGAGGAAGCCAAGGCCTATCAAC | |
| **NKF260** | AAAAGTTCTTCTCCTTTGCTGAAATCAGAGCCGGAAAGCCGTTGG | |
| **NKF261** | GGCTTTCCGGCTCTGATTTCAGCAAAGGAGAAGAACTTTTCACTGG | |
| **NKF262** | TCAATTGCGATTGGCCGGTTTTACTCGAGTTTGTAGAGCTCATCCATG | |
| **NKF263** | AGCTCTACAAACTCGAGTAAAACCGGCCAATCGCAATTGATGTG | |
| **NKF264** | CTGTCAAACATGAGAATTACACCACCCAGCTTGTCGAGCG | |
| **NKF306** | TACCTTCGTGATTACGAAGCATCGTATGGGGCTGACTTCAGGTGC | ***ΔZ6_1479*** |
| **NKF307** | CGCGGCGCTATCGGTGATTGTAATTCTCATGTTTGACAGCTTATCAC | |
| **NKF308** | ACCTGAAGTCAGCCCCATACGATGCTTCGTAATCACGAAGGTATCATG | |
| **NKF309** | AAGTCGCCTTCCCTTGCCGAAGAATTCTAATTTGGCTGGTTTTGACG | |
| **NKF310** | AAACCAGCCAAATTAGAATTCTTCGGCAAGGGAAGGCGACTTC | |
| **NKF311** | AGCTGTCAAACATGAGAATTACAATCACCGATAGCGCCGCG | |
| **NKF312** | CATTAAAGGCTTTTAACGCACGTATGGGGCTGACTTCAGGTGC | ***ΔZ6_0234*** |
| **NKF313** | CCGCTTCCTTGTTATGTCGAGTAATTCTCATGTTTGACAGCTTATCAC | |
| **NKF314** | ACCTGAAGTCAGCCCCATACGTGCGTTAAAAGCCTTTAATGAGGCTG | |
| **NKF315** | ACCCAGTTTACGCCATTCGGCCTTGCCCAATCCTCGTAAAAAAATGATAG | |
| **NKF316** | ACGAGGATTGGGCAAGGCCGAATGGCGTAAACTGGGTCTTG | |
| **NKF317** | CTGTCAAACATGAGAATTACTCGACATAACAAGGAAGCGGTAAGG | |
